# Possible Interference in Protein – Protein interaction as a new approach in micro-inhibition of respiratory pathogens on nasal– oral epithelium: An early on-screen study with reference toSARS-Cov-2–ACE2 binding interference

**DOI:** 10.1101/2021.12.01.470353

**Authors:** Debatosh Datta, Suyash Pant, Devendra Kumar Dhaked, Somasundaram Arumugam, Ravichandiran Velayutham, Pallab Datta

## Abstract

Upper and lower respiratory pathogens – both microbes and viruses –are responsible for very high morbidity, man-hour loss, residual long term clinical conditions and even mortality. In india only, high incidence of annual respiratory infections – both UT and LT – demands prophylactic intervention in addition to all therapeutic interventions available.The issue of respiratory infections is more pronounced now in the backdrop of nearly uncontrolled high incidences of SARS-Cov-2 affection resulting in death and damage of human lives to the extent of hundreds of millions spreading over entire world, with incidence variations from country to country. After the initial unanswered phase of spread of SARS-Cov-2 virus with attendant unseen mortalities, quickest invention of a series of unusual vaccines have stemmed the lethal progress to a very significant extent, although vaccinating each and every human subject – nearly 8 to 9 bn in supremely divided world –economically-- is an unthinkable proposition where economic disparity dictates vaccine availability and implementation.Moreover, being of highly unstable nucleic acid composition, the original virus, by now has a thick set of variants around the globe with variable clinical outcome. Given this complex background of scanty availability and inefficient implementation, there always is a need of a preventive approach which can possibly micro-fix the pathogens, including SARS-2 on nasal epithelium so as to interfere with viral [or any pathogen] entry through specified receptor gate[s] or any other ways. The present formulation is under study -- as a candidate of interference on nasal / oral mucosa for all respiratory pathogens. This brief report describes dry on-screen studies of protein – protein interaction as well as its possible interference by an amino acid Lysine. Phospholipid bilayerresponses in presence of added loads of the same essential amino acid –Lysine – showed unusual and unexplained behavior both in structural integrity as well in spatial orientation.

## Introduction

Two-yr old break of highly contagious and lethal SARS-2 Corona virus and its derivatives along with their rapid spread across the globe with attendant casualties have been one of the most serious challenges to healthcare management and delivery systems anywhere, including in most organized and advanced countries [1-3]. WHO declared the spread and lethality of Covid-19 -- a pandemic in March 2020 with a warning of prolonged struggle with the unstable virus and its derivatives with urgent necessities of preventive, protective and therapeutic agents and formulations [4].

While therapeutic interventions have been put in practice earliest, based on clinical expressions of severe inflammation, uncontrolled intravascular coagulation and multiorgan affections and shutdown, protective approaches by way of variable lines of vaccines -- e.g. mRNA, DNA and whole virus vaccine -- have come into operations with variable undefined and unexplained degrees of protection.

While degrees of protection by available vaccines have been variable, various attendant questions remain unanswered and are being addressed gradually -- globally -- like spacing, number of inoculations, mixing of types, any adjuvant[s] needed, Covid specific antibody titers andtypes with time [sustainability of Ab titer], status of memory cell population, vaccine interactions, responses in immune-compromised and immune-aberrant subjects etc. These coupled with the daunting proposition of vaccinating 8 – 9 billion global population effectively, exhaustively and comprehensively looks like our biggest ever healthcare challenge which may remain unmet ever, given economic, social and educational disparities across the world.

Unfortunately, even a single source of viral repository in future -- anywhere -- will bring possibly few rounds of death and despair that we saw over last two years [which originated officially from one Chinese lady in a Chinese city, finally spreading to the entire world with known lethality].

Given this background, while vaccines will be improved, refined, modified and made more protective and improved therapeutic agents will keep coming with time, there will always be a necessity of an effective preventive approach / formulation / device with proven preventive efficacy, ease of application /use, easy availability, reasonable pricing and nearly zero toxicity / adverse effects.

### Mechanism of anchorage of SARS Covid -2 on respiratory and oral epithelium

Primary respiratory entry of SARS-Covid-2 is through nasal – oral epithelium utilizing homo-trimeric glycoprotein [SPIKE protein S, comprising of S1 and a S2 subunits in each spike monomer] binding to functional receptor human ACE2 [hACE2] and this is a shared mechanism used by earlier SARS–CoV members of Corona family as well as the new SARS-Cov-2 [4]. Binding of the Spike Protein of both SARS–CoV and SARS–CoV-2to the peptidase domain [PD] of ACE2 initiates endocytosis and translocation of both the virus and the ACE2 receptor into the endosomes within the respiratory lining cells. Structure of ACE2in complex with SARS-CoV-2 spike receptor binding domain [RBD] reveals the major RBD interaction regions to be between Q24 – Q42, K353 – R357 and L79 – Y83 of helix H1, a loop in a beta-sheet and the helix H2 respectively [5]. Binding interface consists of 20 residues of ACE2interacting with 17 residues of RBD [5]. Although both ACE2 and Spike protein are heavily glycosylated with many glycosylation sites close to the binding interface, yet the focus has been on amino acid interactions in the ACE2 – Spike protein binding rather than on degree and extent of glycosylation [5]. The main N-glycosylation site on ACE2 in the vicinity of RBD – ACE2 complex which affects binding affinity and viral infectivity has been identified on N90 [6] although there have been 7 N-glycosylation and several O-glycosylation locations on the extracellular domain of ACE2 receptor [5, 6, 8].

### Lysine as a natural interference molecule

This approach of interference in Protein – Protein interaction [between RBDs of incoming pathogens as represented here by SARS-COV-2 and SARS-COV and ACE-2] looks at the possibility of the natural basic amino acid causing interference by binding to the common denominator – ACE2 in a concentration dependent way, where concentration of added Lysine in dry experiments having been considered based on the number of moieties engaged in a particular experiment set vis-à-vis number of Phospholipid moieties allowed in the given set for membrane stabilization.

#### Calculation of concentration of added Lysine

12 Lysine moieties [irrespective of isomeric form] in a phospholipid count of 288 with free water molecules allowed for stabilization of the bilayer gives an aminoacid concentration of 4% [12 /300]. Variations in added amino acid concentration was adjusted in the given way [Lysine 1 – 6 in Fig II, essentially indicates respective percentage concentrations].

**Fig I:**
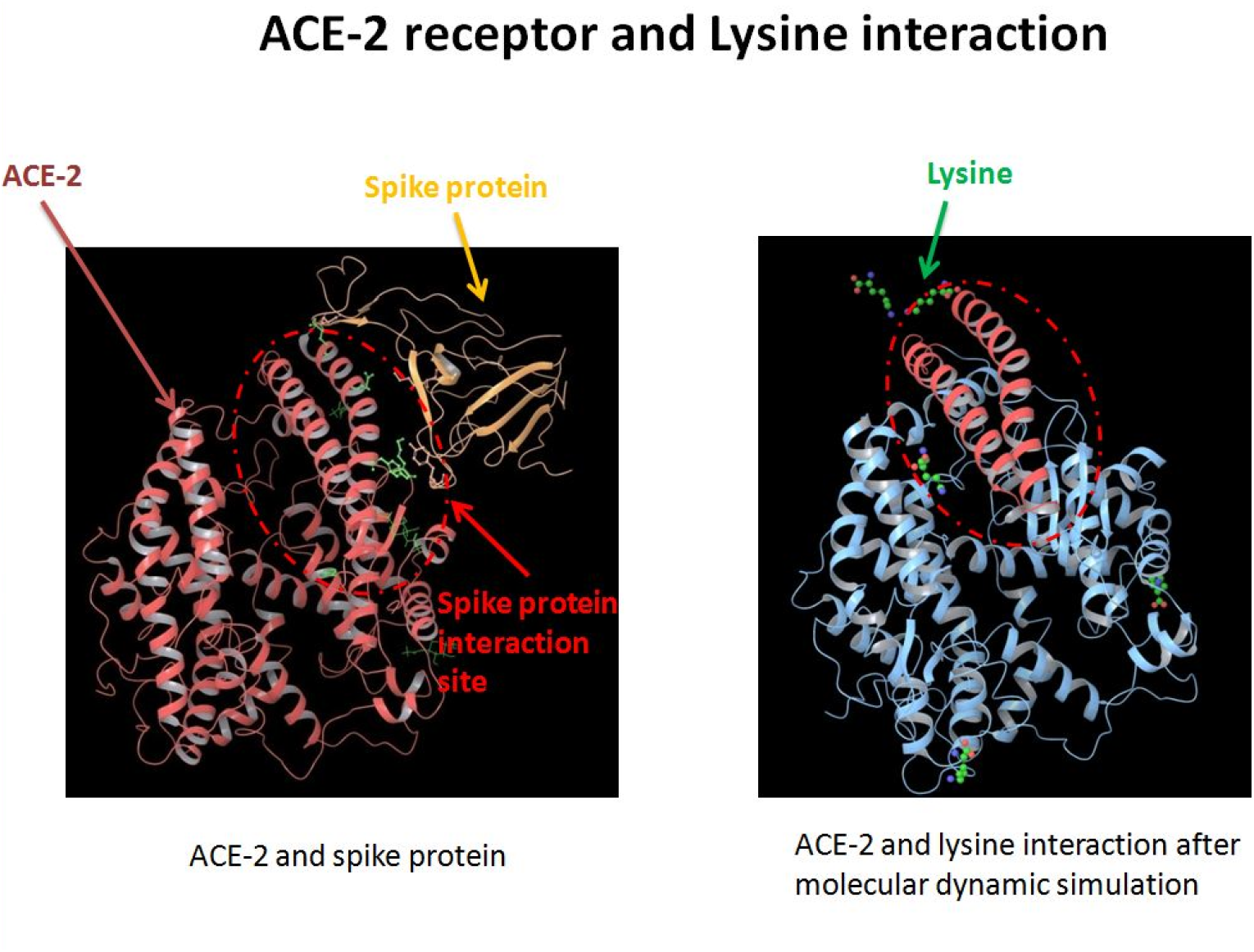
Incremental added Lysine [1-6%] binding to ACE-2 in a [SARS-COV-2 -- ACE2] binding model: Addition of external loads of Lysine was studied in the RBD – ACE2 complex with incremental load being used [As shown]

**Fig II.**
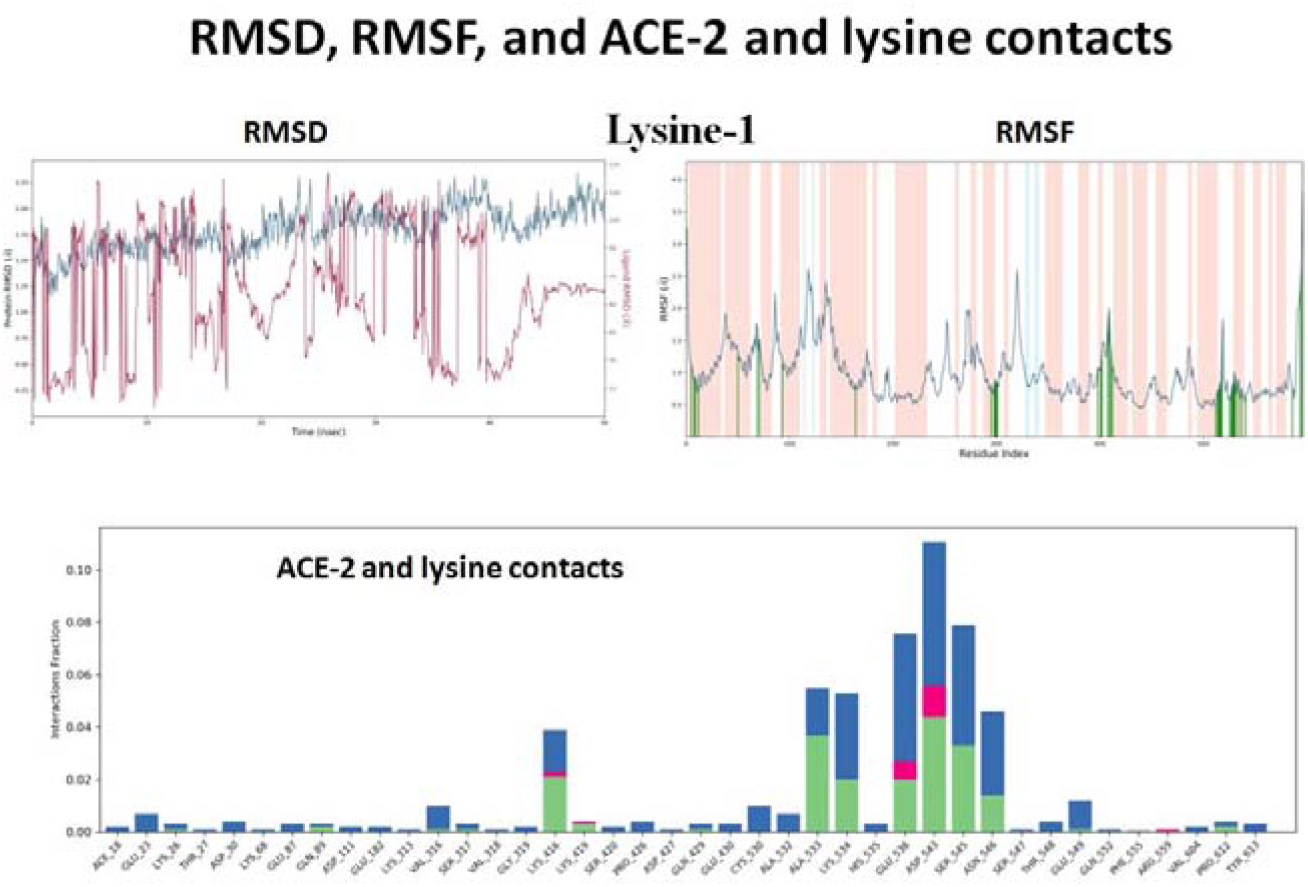
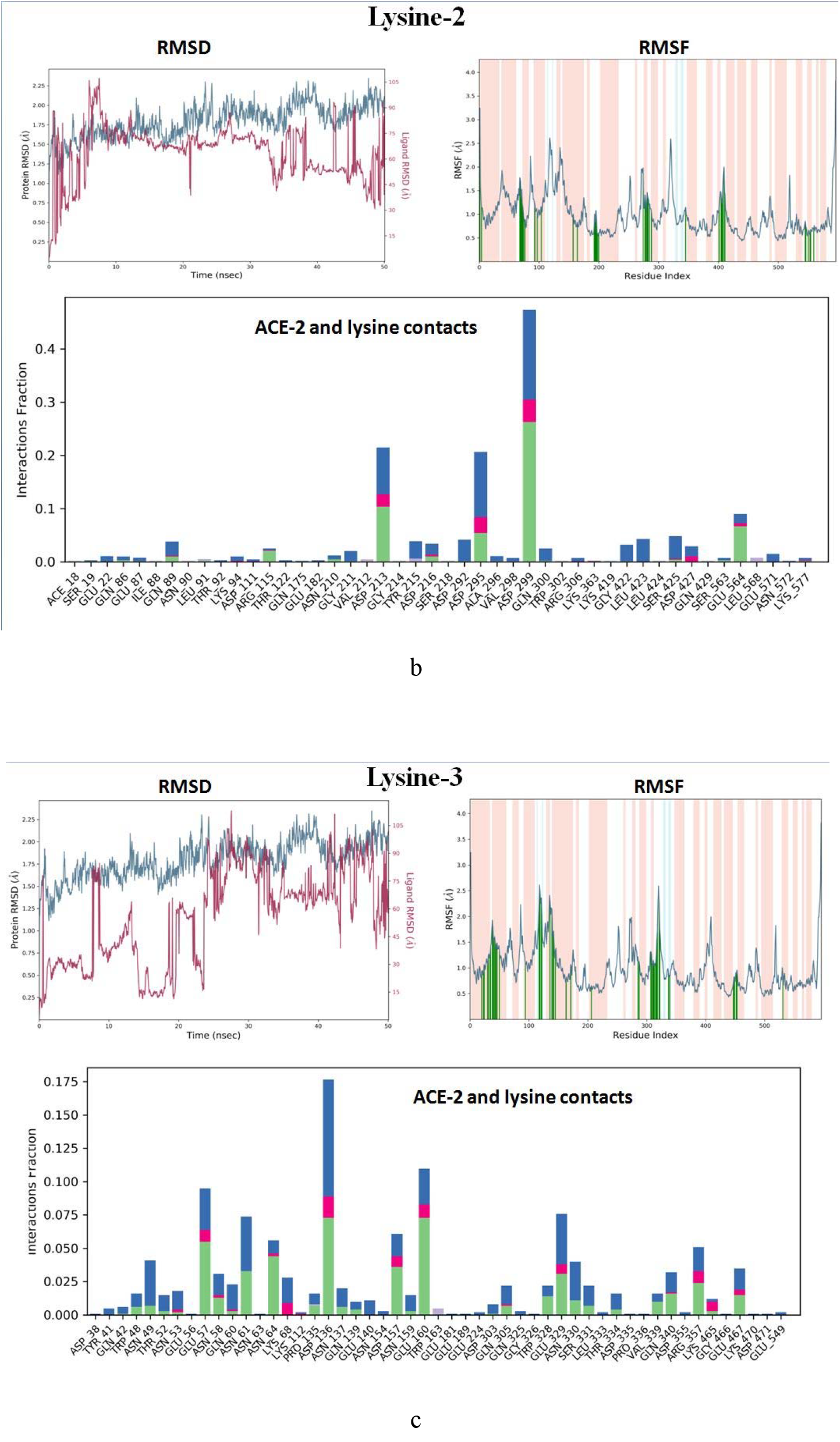
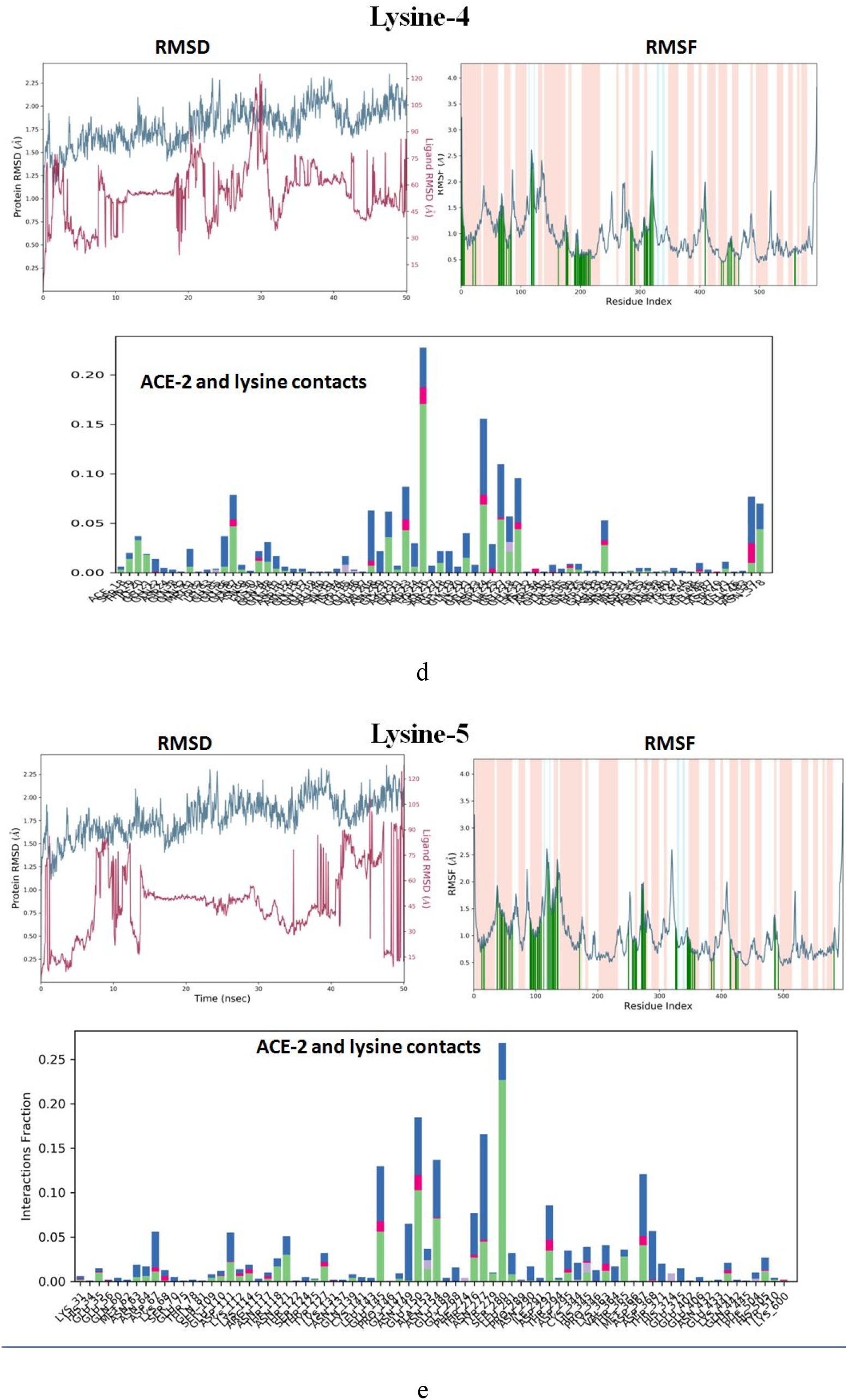
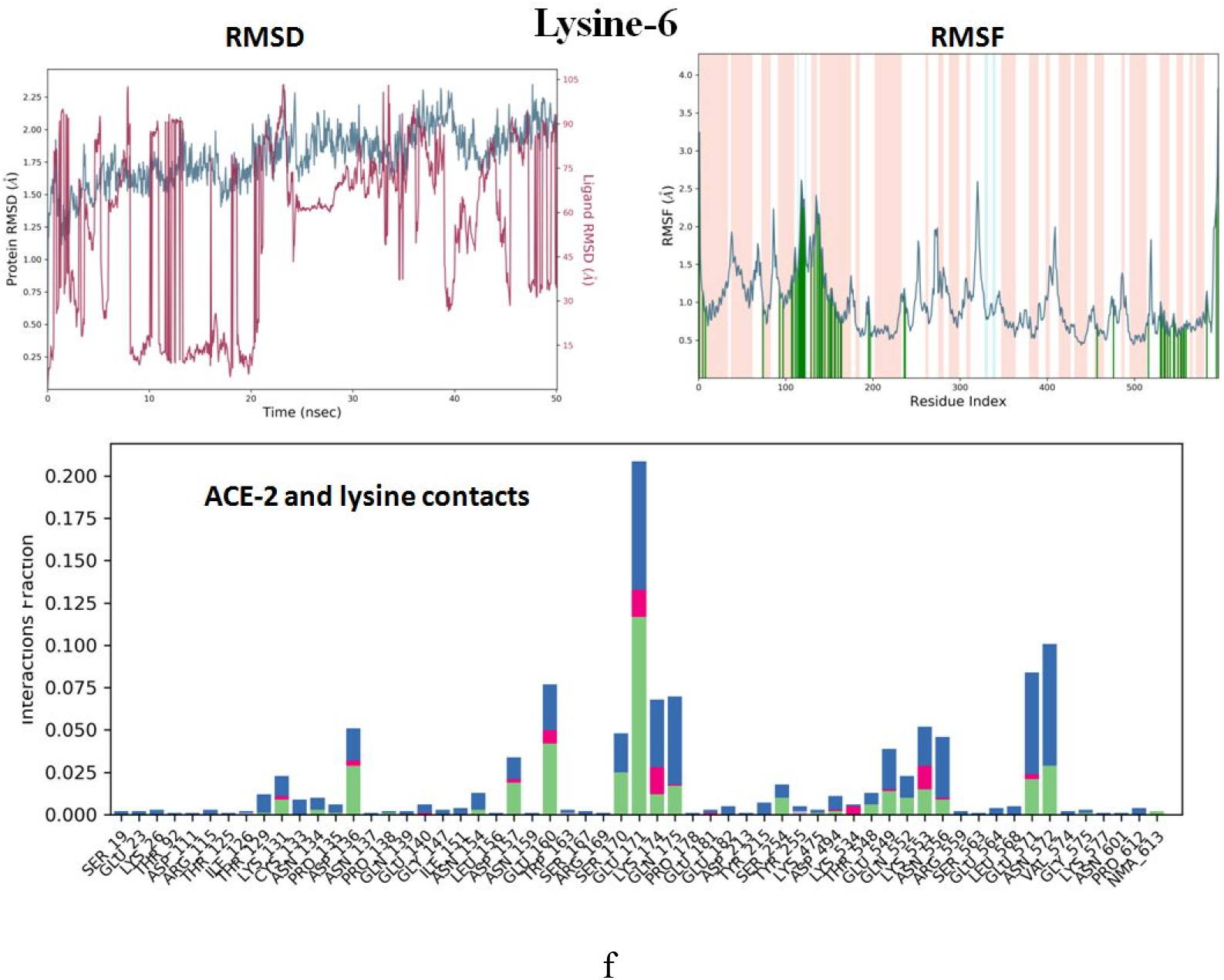
[a – f] -- Interference in Protein – Protein interactions. Lysine binding locations vis-à-vis patterns of interactions with varied loads of Lysine in Protein – Protein binding interferences [Bonding types are expressed in a plot of *interaction fraction* versus* number of residues*, with molecular deviations [RMSD] and fluctuations [RMSF] given above. Green Bar denotes Hydrogen bonding with amino acid R groups; Blue Bar represents stabilizing hydrogen bonding with free water molecule in the vicinity and Pink Bar represents ionic interactions between given Lysine and side chains of amino acids in the exposed segments of ACE2 peptides.

## Rationale

Lysine binding to ACE-2 was studied as a common denominator keeping in consideration allrespiratory pathogens including corona family members [which cause about 30% incidences of Upper and Lower respiratory tract infections in India and south Asia], where pathogen internalization takes place through protein-protein interactions. In addition, prior corona outbreaks in the form of SARS-COV and MERS-COV demand a possible interference model, as possible representatives of other respiratory pathogens. Lysine -monomeric [and oligomeric] - as interference molecule[s] / molecular traps on nasal and oral epithelium need further examination for other respiratory pathogens although a distant parallel may possibly be anticipatedfrom the present interference model. In fact examination of binding domain of SARS-COV with ACE-2 was done with obvious objective of finding any binding pattern and binding homology / overlap between COV-2 and COV [Table I and Figure II a,b,c,d,e,f].

**Table I:**
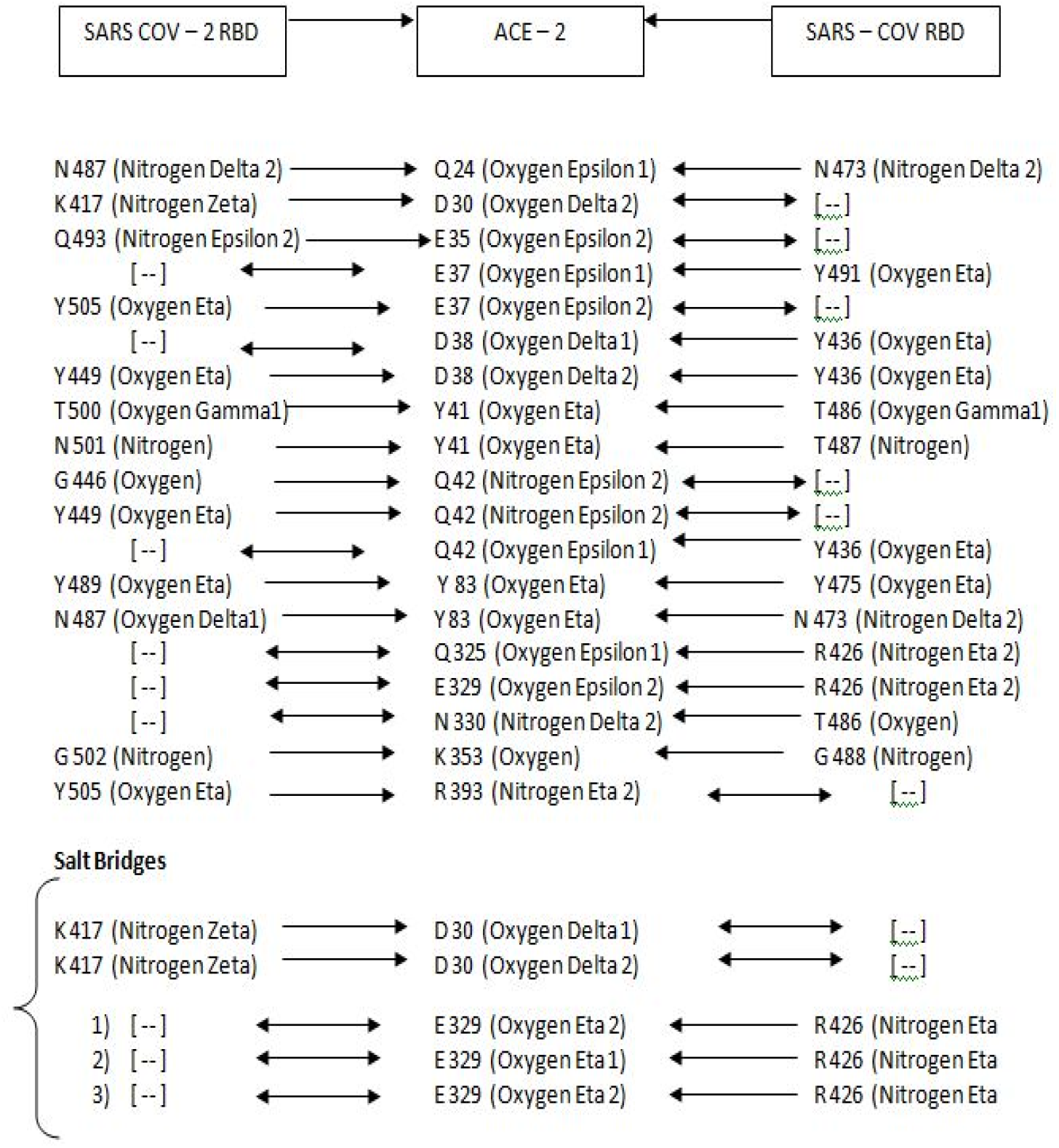
Hydrogen bonds and salt-bridges at RBDs-ACE2 interfaces [4]: Amino Acid interactions [SARS-CoV-2 → ACE2 ← SARS-CoV]:

### Binding stabilities need additional probe

## Results and Discussion

With incremental load [1 – 6%], Lysine – ACE2 binding showed a definite pattern: [Green – Hydrogen Bonding, Pink – Ionic Interactions, Blue – stabilizing water Bonding]

1. Preferential binding locations were obvious as seen in VEGF-Lysine binding patterns [Fig. III; Ref 7]
2. Preferential binding locations were distributed – A533, K534, E536, D543, S545B546 [Fig III, 1%]
3. Nonspecific distributed residual hydrogen bonding and stabilizing water bonding were evident [1%]
4. Nonspecific distributed bonding was evident in 2% exposure over large number of residues with definite shift in preferential binding residues [D213, D295 and D299], with preferential binding residues shifting more upstream [N-Terminal]
5. With given Lysine concentration increasing to 3%, the preferential lysine binding residues shift still further upstream, possibly indicating an unwinding of ACE2 loop with incremental loads of N-acceptors [N49, E57, N58-K68, D136, N137-E160, W328-T334, V339-E467]
6. With further increase in concentration of added Lysine [4%], pattern of upstream shift of nonspecific and specific bindings remains same [between residues 208 and 231 for preferential bindings whereas nonspecific bindings stretched between residues 19 and 101]. Possible reason of upstream bindings remaining same, variation in preferential binding locations between 3% and 4% were between residues 49 – 160 and 208 -231 respectively.
7. Important to note - that 3% added Lysine gives maximum binding overlap of ACE2 residues [49 -160] with the ACE2 residues making active binding site for both SARS-COV-2 and SARS-COV [mainly stretching between 24 - 83 residues and partly between 325 and 329 residues, Table I]. With N-acceptor group load increasing from 1% to 3%, upstream shifts in binding patterns are more likely to interfere with RBD binding domain of ACE2.
8. With further increase in added lysine load, preferential binding locations shift little downstream [between residues 145 and 368]. Although not in the immediate vicinity of RBD binding domains of ACE2, 5% concentration can also possibly induce required conformational changes in the binding domains of ACE2 and this needs experimental substantiation at cell culture, animal as well as in human evaluations. Further increment to 6% load doesnot add any further increments – qualitatively, while in fact gives lesser quantitative adherence.
9. Although approximated in two SARS-Covid RBDs here, parallel binding interferences by lysine and or related molecules can logically be examined and/or extrapolated for otherpossible respiratorypathogens, both in nasal and oral routes of entry, and need be addressed with such binding interferences between any respiratory pathogen and its identified molecular receptor.
10. Bilayer phospholipid interactions with added lysine gave inconsistent responses in lower concentrations -- ranging between shifting membrane discontinuities with time represented here in a scale of 100 ns and membrane deformation in higher concentrations [Fig IV, a - b]. These unexplained responses need further studies simply because of the fact that – if reproducible – it holds the prospect of the natural amino acid -- in variable concentrations – can be used as a possible anti-microbial and anti-viral agent in-vivo [study under progress].

**Fig III:**
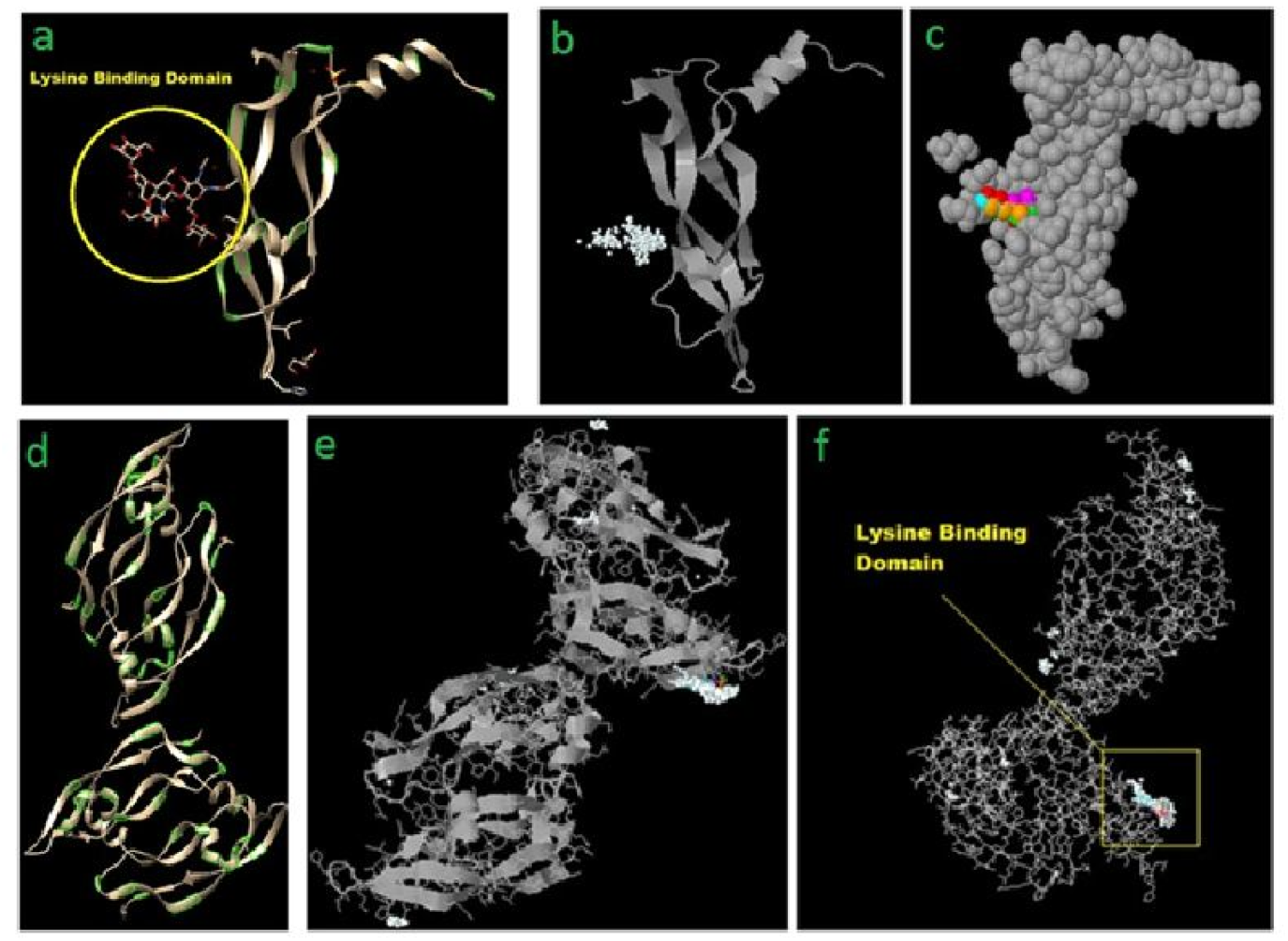
Preferential lysine binding sites in VEGF [7] as a parallel pattern as seen in ACE2 [Fig II, a—f]: Significance of such preferred binding site for a natural amino acid in added loads – much beyond metabolic concentrations -- on such well conserved proteins engaged in vascular reorganizations [controlled angiogenesis] and vaso-motor dynamism remains unclear.

**Fig IV:**
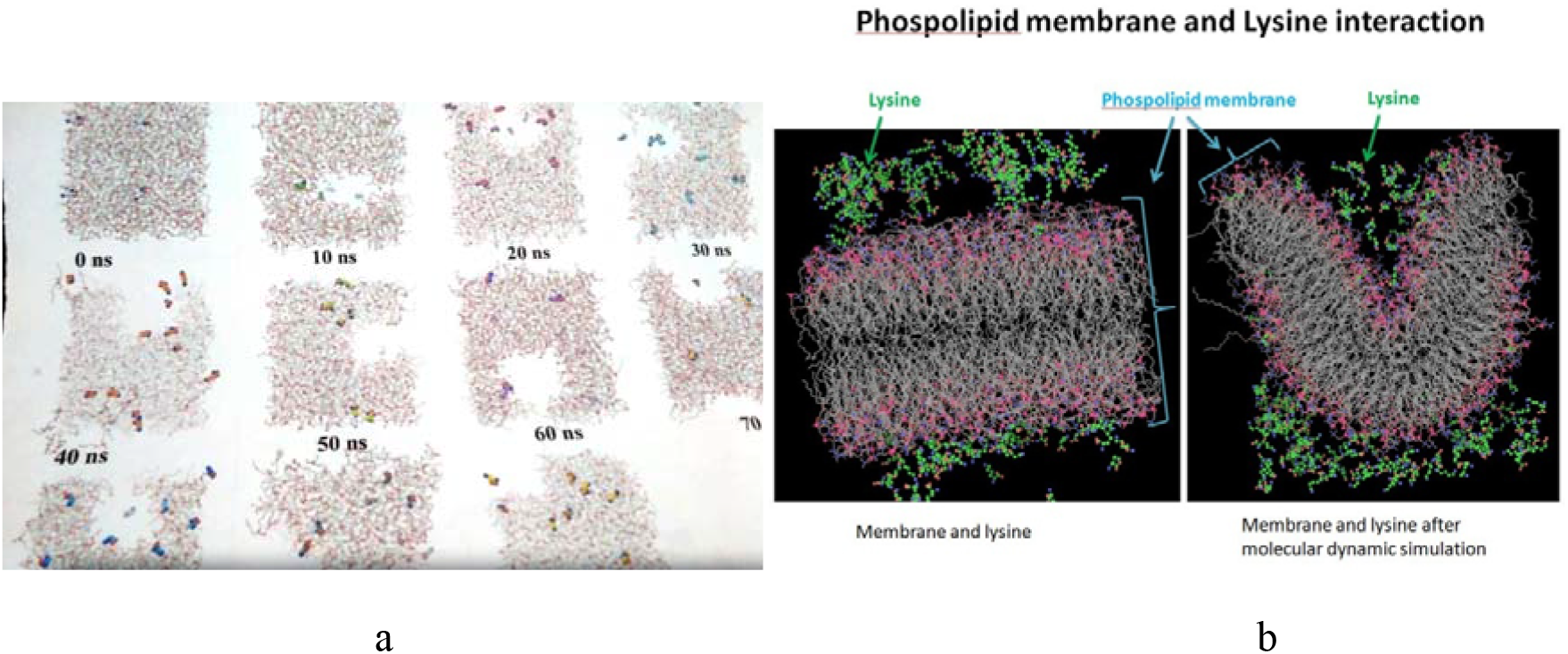
Phospholipid – added lysine interaction and responses were liner with added lysine load where responses varied between unexplained transient travelling and re-coalescing membrane discontinuities to membrane deformation [a, b].

## Conclusion

Bonding analysis of RBDs in SARS-COVID-2 and SARS-COV with ACE2 gives an overall pattern of a concentrated number of H-bondings involving upstream residues between Q24 and Q42 [12 H-bondings], Y83 [2 H-bondings], Q325, Q329, N330, K353, R393 [1 each H-bondings] while D30 and distal E329 gave another set of five salt-bridges on ACE2. These multi-attachments give required anchorage between Spike S1 and S2 complex and ACE2. As a parallel of this observation, given lysine binding in lower conc. [1-3%] in the preliminary study gives a more N-terminal bindings [upstream] involving Asp, Ser, Glu and Asn [#7 in Results and Discussion]. Whether relatively downstream binding locations of higher concentrations of added lysine [4 – 5%] [Fig II, d and e] will cause any spatial molecular deformation[s] in ACE2 as to negatively affect its binding abilities with RBDs of pathogen(s) need be examined.

Preferential lysine binding site[s] is not unique for ACE2 and similar preferred and unexplained high affinity lysine binding site was evident in another model --- VEGF – Added Lysine --- as shown. While biological significance of such high-affinity binding site for a natural amino acid to a protein of such profound importance like ACE2 and VEGF remains elusive, this binding was hypothesized to stabilize VEGF—VEGFr binding leading to in-situ expansion of endothelial cells in ischaemic tissue capillaries. In ACE2 –Lysine binding pattern, a possibility of interference possibly emerges. This needs to be examined in other respiratory pathogens – particularly respiratory virus species [influenza, rhino, adeno and other corona members to name a few].

Interference in Protein – Protein bindings as a possible model of a prospective preventive nasal – oral formulation draws from this parallel more than phospholipid – lysine interactions from this early dry study.

Other current approaches of fixing respiratory pathogens in nasal – oral passages based on uses of anti-microbial active ingredients -- molecular iodine, compounded alcohol -- look much more archaic than the current approach where a non-toxic amino acid –even in loading doses -- is being used as a micro-fixing, micro-entrapment agent on nasal-oral epithelium and proposed mechanism presented is as universal as any anti-microbial APIs and looks ideal for chronic uses in all age groups. Anti-microbial molecules mentioned, in contrast, does not possibly qualify for chronic use and across all age groups. Of course, any complimentary role[s] between oxidizing agent[s] and the amino acid in fixing and eliminating respiratory pathogens in-situ need further investigations.

Additional possible significance of the presented molecular interference model in the presented Spike – ACE2 anchorage might be in the observed well-spread random bindings of the interfering molecule – Lysine – with amino acid residuesof ACE2, superimposed with concentrated locational bindings around particular stretches of residues with varying lysine concentrations examined in this preliminary study [1 – 6%] [Fig II, a-f]. Further detailed analyses of spatial and temporal data on lysine bindings to ACE2 residues along with their stabilities need be done, thus possibly making a new and simpler approach of countering rapidly emerging strains with increasing and varied virulence where existing vaccinecoverage, cross-protection and herd immunity aren’t settled issues and making strain specific immunological protection – a distant dream [B1.1.529]

